# Curling of epithelial monolayers reveals coupling between active bending and tissue tension

**DOI:** 10.1101/806455

**Authors:** Jonathan Fouchard, Tom Wyatt, Amsha Proag, Ana Lisica, Nargess Khalilgharibi, Pierre Recho, Magali Suzanne, Alexandre Kabla, Guillaume Charras

## Abstract

Epithelial monolayers are two-dimensional cell sheets which compartmentalise the body and organs of multi-cellular organisms. Their morphogenesis during development or pathology results from patterned endogenous and exogenous forces and their interplay with tissue mechanical properties. In particular, bending of epithelia is thought to results from active torques generated by the polarization of myosin motors along their apico-basal axis. However, the contribution of these out-of-plane forces to morphogenesis remains challenging to evaluate because of the lack of direct mechanical measurement. Here, we use epithelial curling to characterize the out-of-plane mechanics of epithelial monolayers. We find that curls of high curvature form spontaneously at the free edge of epithelial monolayers devoid of substrate *in vivo* and *in vitro*. Curling originates from an enrichment of myosin in the basal domain that generates an active spontaneous curvature. By measuring the force necessary to flatten curls, we can then estimate the active torques and the bending modulus of the tissue. Finally, we show that the extent of curling is controlled by the interplay between in-plane and out-of-plane stresses in the monolayer. Such mechanical coupling implies an unexpected role for in-plane stresses in shaping epithelia during morphogenesis.

## Introduction

At all scales, from the cell plasma membrane to the petals of flowers, living systems use the ability of thin sheets to form curved shapes adapted to their functions (1, 2). At the tissue scale, epithelial monolayers bend and fold as part of developmental morphogenesis. This process requires the regulated activity of biological actors to generate spatially patterned physical forces. So far, folds formation in epithelial monolayers on the minute time-scale has been shown to result from local contraction of the actomyosin cytoskeleton. In particular, bending moments arise either through the formation of apico-basal acto-myosin cables (3, 4) or through constrictions caused by an asymmetric distribution of Myosin II along the apico-basal axis (5, 6). This latter process drives tissue morphogenesis during development, for instance during the mesoderm invagination in *Drosophila* (7–9), but can also accompany disruption of epithelial architecture in disease, for example during tumor growth in the pancreatic duct (10). Epithelial folding can also occur in response to in-plane compressive deformations, when their magnitude is sufficient to dissipate tissue pre-tension generated by Myosin contractility (11). Although each of these mechanisms in isolation can give rise to folding, theoretical and experimental insights suggest that variations of in-plane stresses at the tissue-scale can act in parallel to apical or basal constrictions to define the three-dimensional shape of epithelia (12–14). Yet, in contrast to the mechanical properties and active forces generated by epithelial monolayers within the plane (11, 15–20), out-of-plane forces and mechanical properties of epithelia (active torques and bending modulus) have not been characterized. Consequently, the relative magnitudes of forces acting in-plane and out-of-plane remain unknown, making the contribution of out-of-plane stresses to morphogenesis challenging to assess.

Here, we use epithelial curling to characterize out-of-plane tissue mechanical properties. Curling is a mechanical phenomenon which occurs at the free edge of elastic materials endowed with a spontaneous curvature. For example, curling has been observed in the cell membrane during malaria parasite egress (21) or on a sheet of paper swelling at a liquid surface (22). In this article, we find that curling occurs at the free edge of epithelial monolayers devoid of substrate *in vivo* during *Drosophila* leg eversion and *in vitro* in suspended Madin-Darby Canine Kidney (MDCK) monolayers. We show that the high curvature of the curls is controlled by the asymmetric localization of Myosin II motors along the apico-basal axis. By unfurling the tissue via micromanipulation, we measure the amplitude of the resulting active torques as well as the bending modulus of the monolayer. We find that the bending energy of the monolayer is comparable to its in-plane elastic energy. This property induces a coupling between the in-plane and out-of-plane tissue shape, which can be understood quantitatively by modeling the tissue as a continuous thin sheet endowed with a spontaneous curvature. Altogether, these findings show that high spontaneous curvature generated by an apico-basal anisotropy of contractility allow epithelial monolayers to regulate their three-dimensional shapes according to dynamic in-plane boundary conditions.

## Results

### Epithelial monolayer curling depends on myosin II polarization

To investigate the importance of out-of-plane forces in shaping epithelia, we examined curling in monolayers. We hypothesised that epithelial tissues possessing an apico-basal asymmetry in myosin distribution along with a free edge and negligible matrix foundation should curl. Suspended epithelial monolayers meet these conditions. Indeed, these tissues are generated by culturing the epithelium on a scaffold of type I collagen, which is polymerized between two parallel glass plates and is then removed via enzymatic digestion (Fig 1A, see Methods) (15, 23). Upon matrix removal, we noticed that the free edge of MDCK monolayers retracted and took on a paraboloid shape (N=14/14, Fig 1A & S1A, Movie 1). These shape changes in the plane of the tissue were accompanied by shape changes in the transverse direction (yz-plane) where the tissue curled towards its basal side (Fig 1A & 1B, Movie 2). We extracted the radius of curvature of the curls *R*_*c*_ = (12.7 ± 1.1)μm (see Fig S1B & S1C for method) which was comparable to the monolayer thickness *h* = (18.6 ± 1.3)μm. Together with the lack of interstice observable within the curl (Fig S1D), this shows that MDCK epithelial monolayers curl as much as volume exclusion and steric interactions allow. We thus hypothesized that such high curvature stems from torques actively generated by an apico-basal asymmetry in the distribution of Myosin II molecular motors. To test this hypothesis, we imaged the distribution of GFP-tagged Myosin II-A and -B in substrate-free MDCK monolayers. Both myosins were enriched at the basal surface and lateral junctions compared to the apical surface (Fig 1C, 1D & S1E). This asymmetry of contractility along the apico-basal axis is consistent with the orientation of the curl towards the basal side of the epithelium. To directly prove that active forces were at the origin of tissue curling, we treated monolayers with Y27632, an inhibitor of Rho-kinase. In these conditions, the tissues showed a reduced asymmetric localisation of myosin II, especially myosin II-B (Fig 1D, 1E & S1F). Concomitantly, the curvature at the tip of the tissue was reduced 2-fold (Fig 1F-1H).

**Fig. 1.**
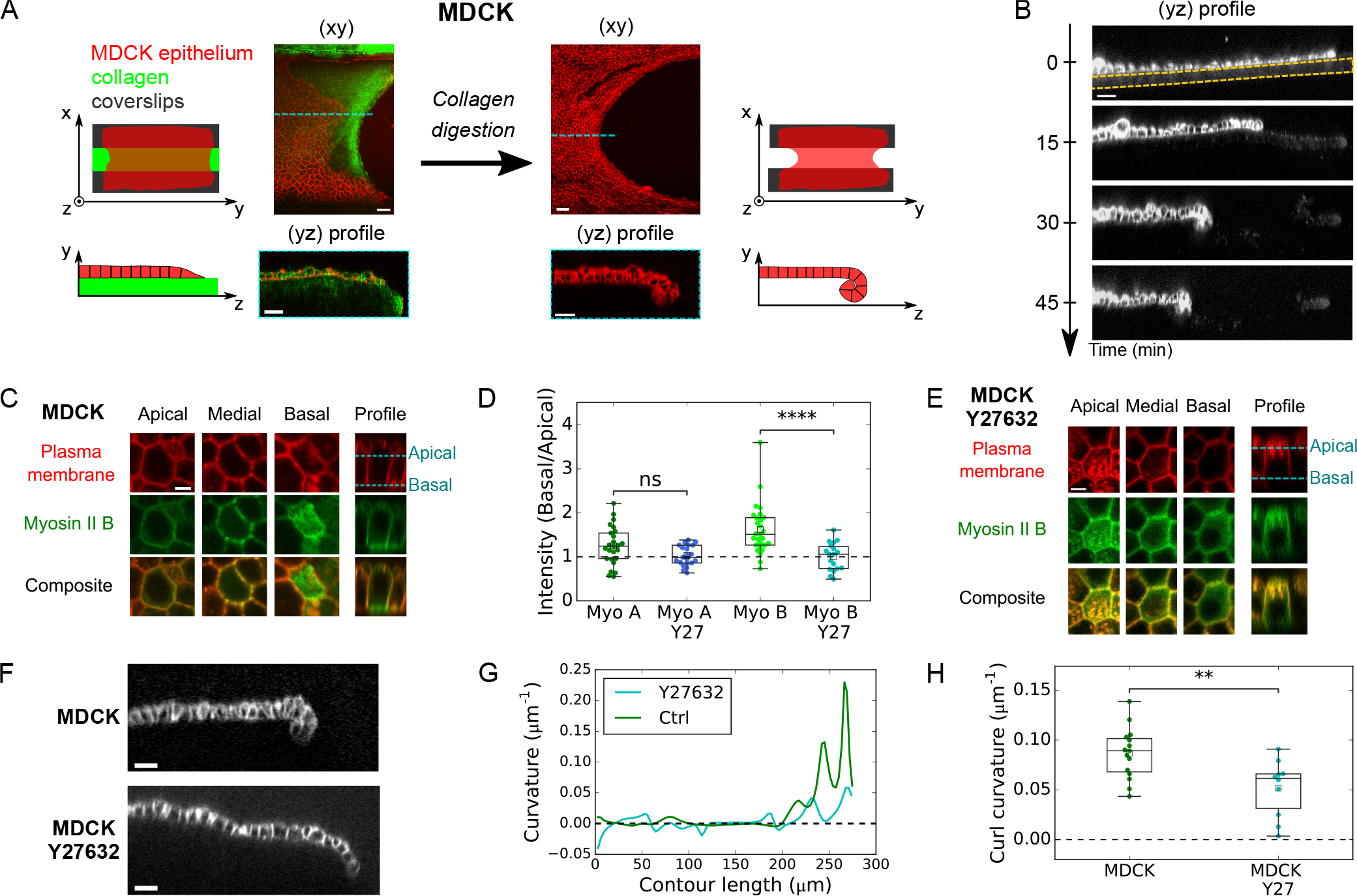
Curling at the free edge of epithelial monolayers is controlled by asymmetry in Myosin II across tissue thickness. (A) Effect of substrate digestion on the free edge of an epithelial monolayer. Left: An MDCK monolayer (red) grows on a device composed of two parallel coverslips connected by a collagen bridge (green). Right: After collagen digestion, the monolayer has retracted in the xy-plane and curled in the yz-plane. (B) Profile view (yz-plane) of an MDCK monolayer as digestion of the collagen substrate proceeds. Collagenase is introduced in the medium at t = 0min. Cell membrane is marked with CellMask. The initial thickness of the collagen substrate is indicated by the dashed yellow line. Scale bar: 30μm. (C) Localisation of Myosin II B-GFP in the apical, medial and basal regions of an MDCK substrate-free monolayer. Plasma membrane is marked with CellMask (red). Scale bar: 5μm. (D) Ratio of Myosin II-GFP average fluorescence intensity in the basal surface to the apical surface (see method in Supplementary Information). Myo A: Myosin II-A (N=25 cells), Myo B: Myosin II-B (N=27), Myo A Y27: Myosin II-A treated with 25μM Y27632 (N=21), Myo B Y27: Myosin II-B treated with 25μM Y27632 (N=18). Data extracted from N=3 monolayers minimum for each condition. (E) Localisation of Myosin II B-GFP in the apical, medial and basal regions of an MDCK substrate-free monolayer treated with 25 μM Y27632. Plasma membrane is marked with CellMask (red). Scale bar: 5μm. (F) Profile view (yz-plane) of the free edge of a suspended MDCK monolayer in untreated conditions (top) and after treatment with 25 μM Y27632 (bottom). Cell junctions are marked with E-Cadherin-GFP. Scale bars: 20μm. (G) Local curvature along contour length of the monolayers shown in F. (H) Boxplot of curl curvature, measured at the monolayer free-edge in untreated (N=15) and Y27632 treated tissues (N=10).

To establish whether epithelial curling occurs *in vivo*, we searched for an *in vivo* tissue which possesses a free edge and detaches from its extra-cellular matrix. The peripodial epithelium enclosing the *Drosophila* leg imaginal disc is known to meet these conditions. During early metamorphosis (24), the leg imaginal disc turns itself inside out (everts) (25, 26). To allow eversion, the peripodial epithelium first detaches from its basal extracellular-matrix before rupturing, which generates a free edge that expands through myosin II-dependent retraction (27) (Fig S2, Movie 3). To focus on the mechanism of retraction at the cell scale, we recorded cell shape changes during peripodial retraction. Together with large reduction of cell area, the peripodial epithelium curled (N=9/13) towards the basal side (N=9/9) (Fig 2A & 2B, Movie 4), suggesting that the basal surface of the epithelium is more contractile than the apical surface. This was consistent with enrichment at the basal domain of Sqh, the homolog of myosin II in *Drosophila*, and the magnitude of the asymmetry was comparable to what we observed in suspended monolayers (Fig 2C & 2D). Accordingly, the radius of curvature *R*_*c*_ of the peripodial epithelium was close to its thickness, as found in MDCK suspended monolayers (Fig 2E). These results show that, in the absence of substrate interference, a myosin II asymmetry along the apico-basal axis can control the out-of-plane curvature at the free edge of epithelial monolayers *in vivo* and *in vitro*.

**Fig. 2.**
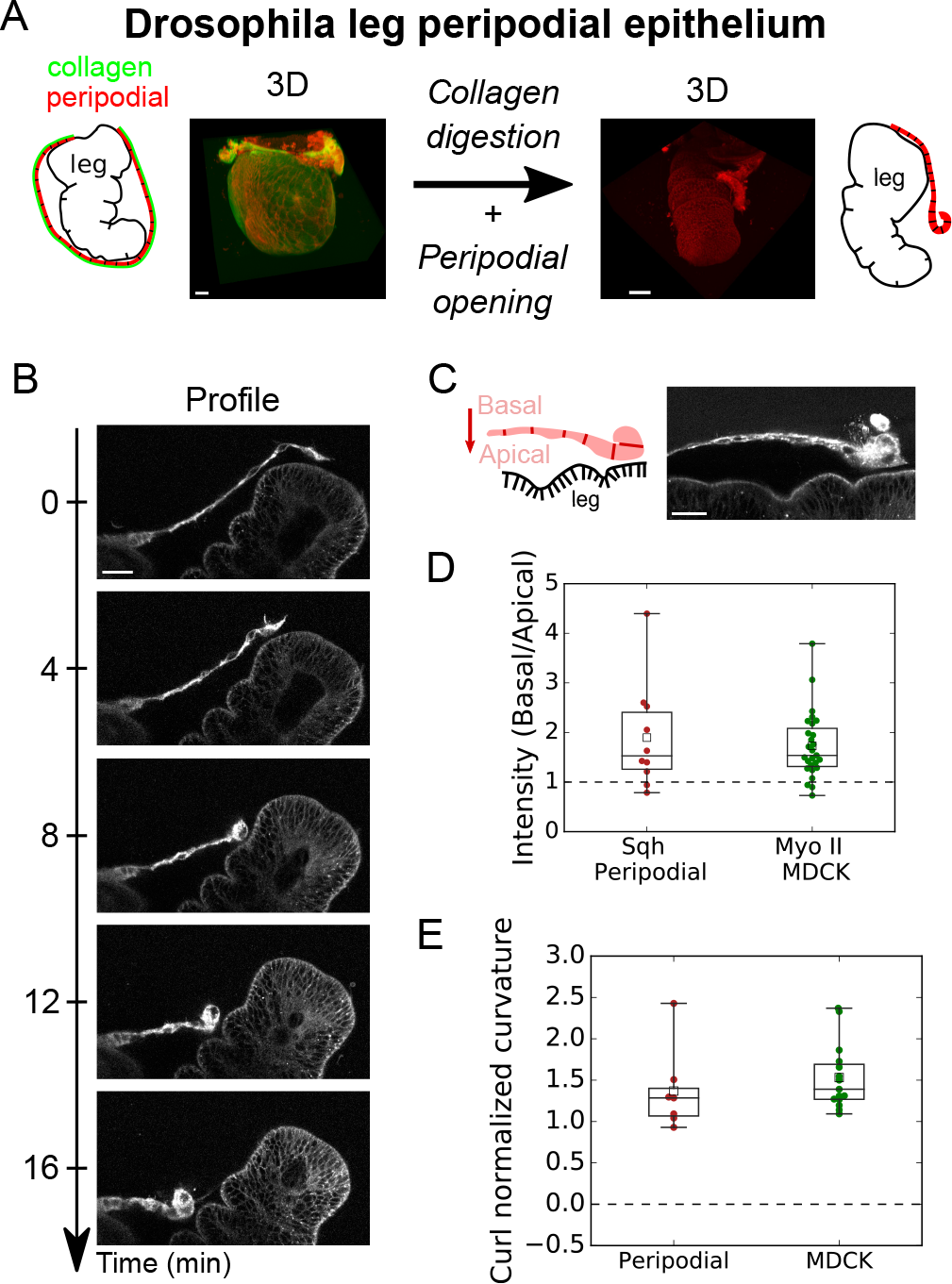
The peripodial epithelium curls in vivo during *Drosophila* leg disc eversion. (A) Fate of peripodial epithelium during *Drosophila* leg disc eversion. Before eversion (left), the leg epithelium is encapsulated in the squamous peripodial epithelium whose basement membrane (Vkg-GFP, green) is intact. After collagen digestion, the peripodial epithelium ruptures and withdraws (right) allowing leg eversion. Cell membranes are marked with CellMask (red). Scale bars: 20μm. (B) Time series of a retracting peripodial epithelium imaged with confocal microscopy. Profile view in the plane perpendicular to the retraction front. Cell membranes are marked with CellMask (red). Scale bar : 20μm. (C) Distribution of Sqh-RFP (Myosin II homolog) during Drosophila leg eversion. Scale bar: 20μm. (D) Ratio of average fluorescence intensity in the basal surface to the apical surface (see method in Supplementary Information). Intensity of Sqh-RFP for the peripodial epithelium (N=10) and Myosin II-GFP (pooled for Myosin II-A and II-B, N=52 cells from 9 monolayers) for MDCK suspended monolayers. (E) Boxplot of normalized curvature (ratio of average tissue height *h* over average radius of curvature *R*_*c*_) at the monolayer free edge in the peripodial epithelium (N=7) and in MDCK monolayers (N=15).

### Characterisation of active torques and the bending modulus

We then aimed to directly test if curls were generated by active torques by measuring the force necessary to unfurl MDCK epithelia. To do this, we brought a glass needle of stiffness *k* into contact with the curled region of a monolayer and imposed a displacement *D*(*t*) at its base via a motorized micromanipulator (Methods). Simultaneously, we imaged the change of curvature of tissue in the yz-plane and measured the deflection δ(*t*) of the cantilever using con-focal microscopy (Fig 3A, 3B & S3A, Movie 5). The force required to unfurl the tissue *F*(*t*) = *k*δ(*t*) was on the order of tens of nN (Fig S3B). Confocal stacks of the tissue before and after force application were acquired to determine the width wc of the region affected by unfurling, we acquired (Fig 3C). Then, the approximate stress required to unfurl one third of the curl length 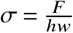 (where *h* is the thickness of the tissue) was found to be on the order of 30 Pa (Fig 3D).

**Fig. 3.**
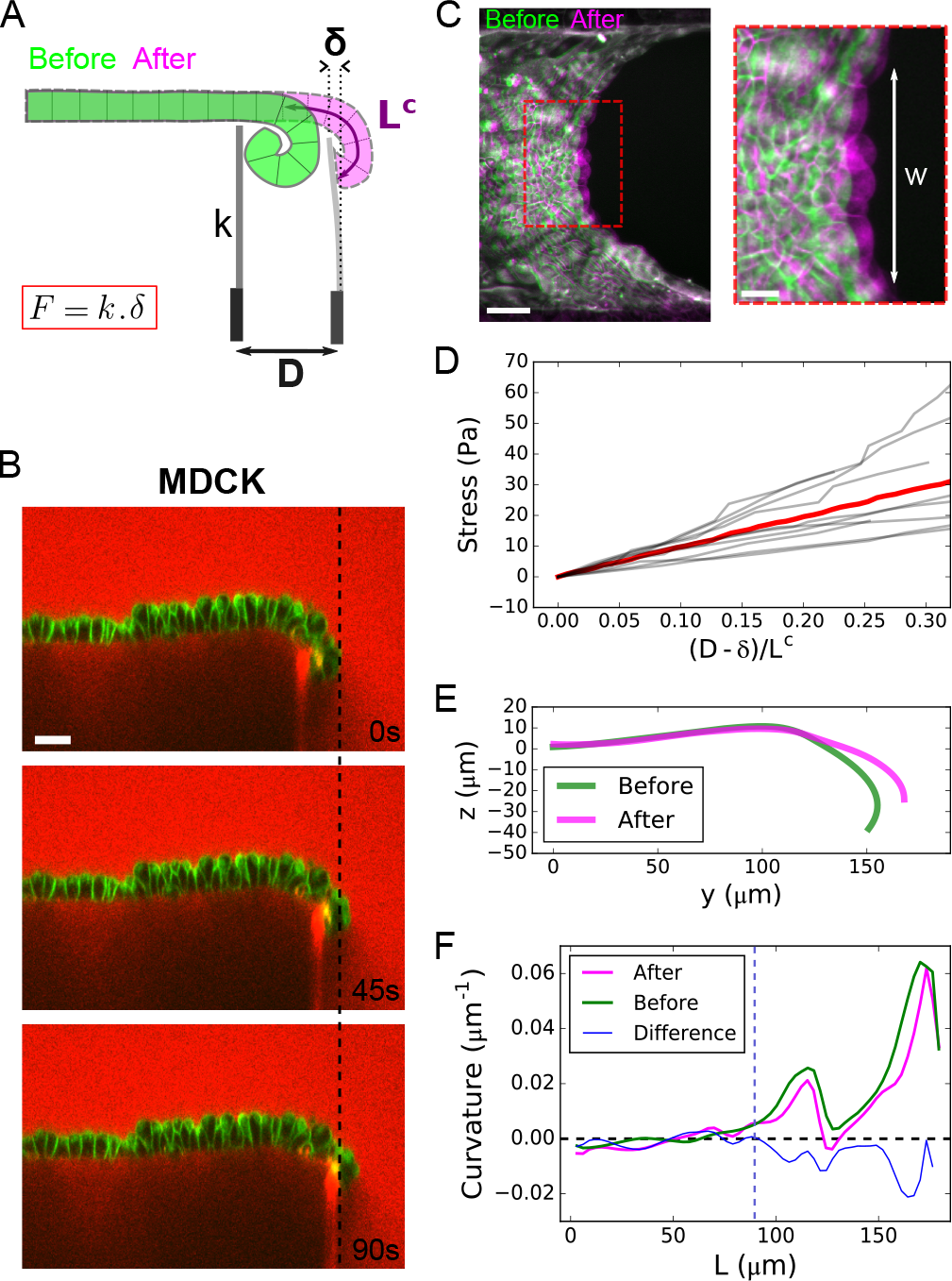
Characterisation of active torques and bending modulus of epithelial mono-layers. (A) Diagram of the setup to measure the bending modulus of an epithelial monolayer. A flexible glass capillary of stifness *k*, serving as a force cantilever, is approached close to a tissue curl (green) of curled length *L*^*c*^. The displacement D imposed at the base of the cantilever generates a deflection δ of the cantilever, due to the restoring force of the flattened tissue (pink). (B) Time series of an MDCK monolayer imaged in profile during a ramp of displacement of the force cantilever. Dextran-Alexa647 was added to the medium (red). Note that this dye also stains the tip of the cantilever underneath the monolayer, which facilitates the measurement of its displacement. Cell membranes are marked with CellMask (green). Scale bar: 30μm. (C) Overlay of projected confocal stacks showing the free edge of an MDCK suspended monolayer before (green) and after (magenta) unfurling of the mono-layer by the force cantilever. wc denotes the width of unfurled tissue. Scale bar: 50μm. Red inset shows a zoom on the deformed region of width w. Scale bar: 20μm. (D) Stress variation along a ramp of displacement imposed at the cantilever base. The displacement of the cantilever tip is quantified by the ratio 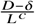, which is equal to 1 when the curl is fully unfurled. The grey lines represent N=9 separate experiments, the red line represents the average. (E) Typical profile of monolayer before (green) and after (pink) unfurling of the tissue by the cantilever. (F) Typical variation of curvature of the monolayer along its contour length L before (green) and after (pink) unfolding by the cantilever. The difference between the two curves is plotted in blue. The dashed blue line indicates the transition between curled and non-curled regions of the tissue corresponding to the position where the blue curve departs from the x-axis.

Because the force applied on the cantilever by the tissue did not relax after unfurling (Fig S3C), we could also use this method to define a bending modulus of the epithelial monolayer. The displacement of the cantilever resulted in a change of curvature of the curled region (Fig 3E & F) along with a small stretch of the bulk of the tissue (Fig S3D). Thus, the work transferred to the tissue via the cantilever displacement is equal to the sum of the variation of bending energy of the curl and stretching energy of the bulk (see details in Appendix 1). This allowed us to estimate a bending modulus of the tissue: *B* = (3.7 ± 0.4).10^−13^ *N* .*m*, close to that predicted by the classical Kirchhoff-Love model of thin plate: *B*_*K L*_ = (2.1 ± 0.2).10^−13^ *N* .*m* and consistent with a recent estimate extracted from buckling of MDCK monolayers due to tissue-growth in long time-scale experiments (28).

### Spontaneous curvature is homogeneous throughout the epithelium

During wound healing, cells at the free edge acquire different characteristics from those in the bulk (29). Therefore, we asked whether curling emerged from specific cell properties at the edge of the tissue or from a spontaneous curvature present everywhere in the bulk of the tissue. For this, we used laser ablation to generate a rectangular cut in the centre of MDCK monolayers (150 × 15 μ*m*^2^, ≃ 16 × 2 cells) whose long axis was oriented parallel to the coverslips (Fig 4A). Ablation of monolayers growing on a collagen substrate was followed by wound healing (Fig S4A, Movie 6), whereas ablation of suspended monolayers induced retraction of the tissue which took on an ellipsoidal shape in the xy-plane (Fig 4A, Movie 7). Simultaneously, the monolayer curled basally in the plane perpendicular to the longest axis of the cut (xz-plane) (Fig 4B, Movie 8). Despite the difference of orientation and location, the curvature at the tip of ablated tissue was similar to curls occurring at the free edge of monolayers after collagen digestion (Fig 4C). The fast time-scale (~ tens of seconds) at which the new free edge could curl indicates that all cells in MDCK monolayers are endowed with a constant isotropic spontaneous curvature which can bend the tissue and lead to curling wherever a free edge is generated. Thus, curling is not a specific property of cells at the tissue edge in our culture conditions. Furthermore, the tissue spontaneous curvature possesses an active origin since Rhokinase inhibition led to the same reduction of curvature at the ablated site as at the tissue boundary (Fig 4C, S4B & S4C, Movie 9).

**Fig. 4.**
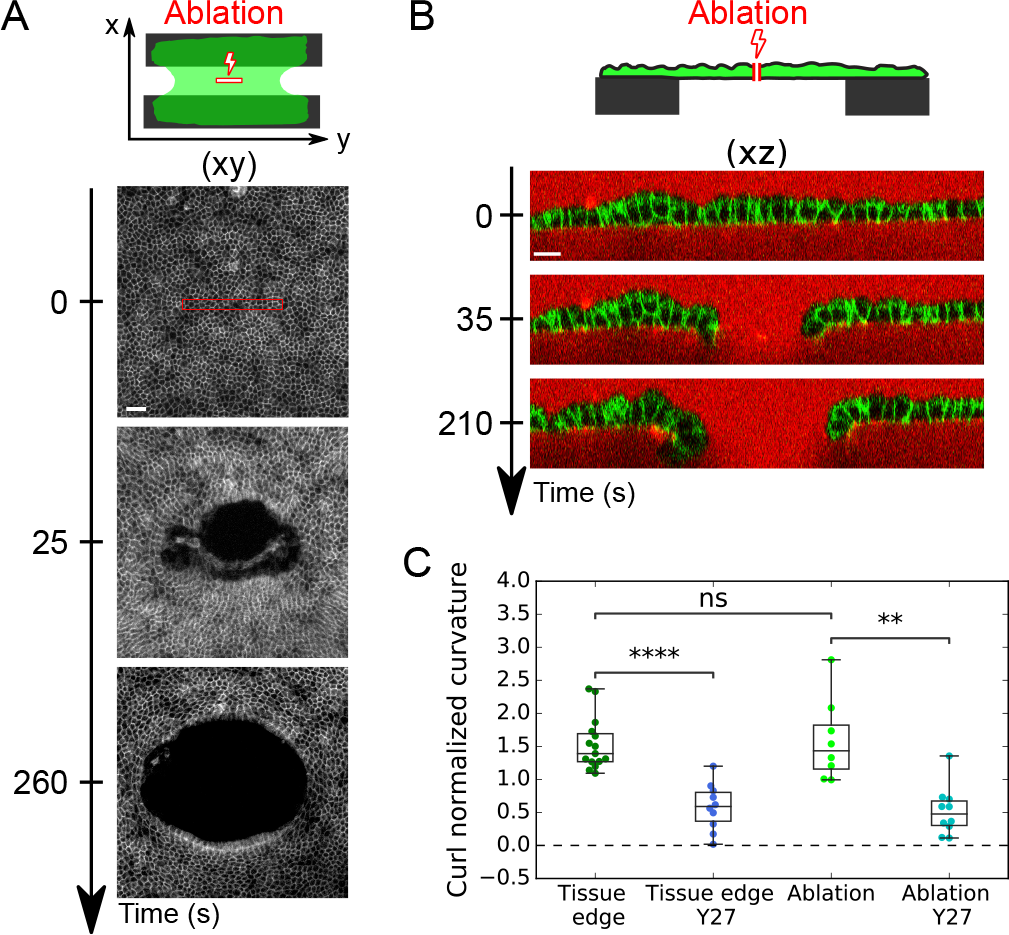
Active spontaneous curvature is a bulk property of the tissue. (A) Time series of the ablation of a suspended MDCK monolayer. Laser ablation was performed inside the red rectangle at time t=0s. Cell junctions are marked with E-Cadherin-GFP. Scale bar: 50μm. (B) Time series of the profile view (xz-plane) of an MDCK suspended monolayer before and after laser ablation. Cell junctions are marked with E-Cadherin-GFP. The medium is marked with Dextran-Alexa647 (red). Scale bar: 15μm. (C) Boxplot of the curl normalized spontaneous curvature (*h*/*R*_*c*_) of suspended MDCK monolayers at the free tissue edge or at the edge created by laser ablation in presence (N=10) or not (N=8) of Y27632 (Y27).

### The extent of curling is controlled by tissue boundaries

Together our results show that myosin II asymmetry controls the spontaneous curvature of the tissue but the parameter limiting the extent of curling at the free edge remains unclear. Notably, in both free edges created in the bulk by ablation and free edges at the tissue boundary, we observed that cells are elongated along the axis tangential to the edge (Fig 5A & 5B). Thus, we hypothesized that curling could be limited by the in-plane tissue tension. Indeed, sheet-like materials with a spontaneous curvature tend to curl naturally to relax their bending energy. This effect deflects the tissue inwards. However, when the sheet is clamped at two of its ends, any deflection of the free edge created by curling will also stretch the free edge. Such stretching could ultimately limit tissue curling because of monolayer elasticity (Fig 5C). In sum, the balance between bending and in-plane elastic energies may define the equilibrium three-dimensional shape of the monolayer. This is in line with the paraboloid shapes observed at the free edge of the tissue in the xy-plane or after laser ablation (Fig 5A). To test this, we first completely abolished tissue tension by cutting monolayers along the full length of the tissue-coverslip interface. After the cut, the tissue curled over its entire length, especially in the xz-plane perpendicular to the cut (Fig 5D, Movie 10), confirming that tissue clamping and the resulting in-plane tension could prevent tissue curling.

**Fig. 5.**
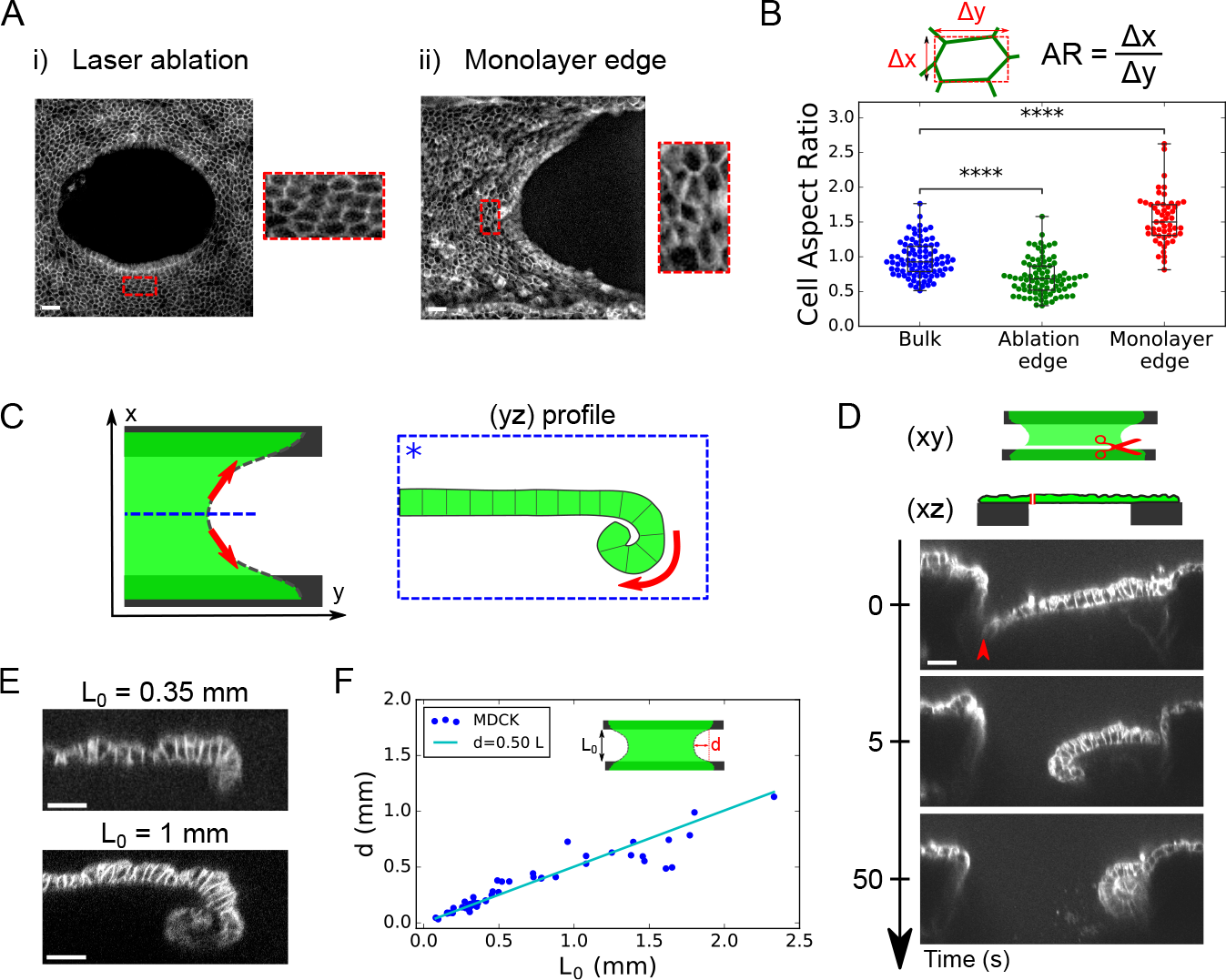
Tissue boundaries control the extent of curling. (A) Maximum projection of confocal stacks showing the free edge of an MDCK suspended monolayer. (i) 260s after laser ablation in the tissue bulk (ii) at the tissue boundary free edge. Insets show stretched cells in the direction tangential to the edge. Cell junctions are marked with E-Cadherin-GFP. Scale bars: 30μm. (B) Boxplot of cell aspect ratio: in the bulk before ablation (blue, N=89 cells), in the vicinity of the ablation site (green, N=87) as in A-(i) and in the vicinity of the free edge (red, N=53) as in A-(ii). N=5 different monolayers in each condition. The aspect ratio (AR) is computed as described in the diagram above the graph. (C) Diagram illustrating the proposed balance of forces in-the-plane and out-of-plane. The monolayer (green) possessing a spontaneous curvature naturally curls at the free edge to relax its bending energy. Curling deflects the tissue inwards (box dashed blue line). Such deflection generates stretching of the free edge (red arrows), which ultimately limits curling because of monolayer elasticity. (D) Time series of a profile view (xz-plane) of an ablated suspended MDCK monolayer (N=12). Ablation was performed along the full length of the tissue at the interface with the coverslip (see diagram). Red arrow-head shows the location of the cut. Scale bar: 30μm. (E) Representative profiles (yz-plane) of MDCK suspended monolayers acquired by confocal microscopy for monolayers of different lengths *L*_0_ (N=15 for *L*_0_ =0.35mm; N=5 for *L*_0_ =1mm). (F) Deflection d of MDCK monolayers free edge as a function of initial length *L*_0_ . The line in cyan shows the linear fit as suggested by our model (see Appendix 2).

### Spontaneous curvature induces a coupling between in-plane and out-of-plane tissue shape

To further assess this idea, we modeled the epithelium as a continuous rectangular elastic sheet endowed with a spontaneous curvature (see Appendix 2). In this framework, the extent of curled tissue at the free edge is a function of the sheet’s initial length *L*_0_, its in-plane deformation 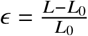 and the mechanical properties of the tissue: the 2D elastic modulus *E*, the bending modulus *B* and the spontaneous curvature *C*_0_. Importantly, all these parameters could be directly measured in our setup. A first prediction of the model, which was confirmed experimentally, is that the deflection *d* (corresponding to the length of tissue that curls at the center of the free edge) increases proportionally to the unstrained tissue length *L*_0_ (Appendix 2, Fig 5E & 5F).

Conversely, our model predicts that, in response to uni-axial stretch, the curl should unfurl as this shape change reduces the length of the free edge and the in-plane elastic energy of the sheet (Fig 6A). To directly reveal such coupling, we dynamically controlled the length of the tissue along the x-axis (Fig S5A) and monitored the effect on the curl length at the monolayer free edge (yz-plane). We first imposed a slow ramp of deformation ranging from +20% to −20% at a strain rate of 0.5%.s^−1^. In this regime, MDCK monolayers behave elastically with stress increasing proportionally to strain (Fig 6B) (11). Here, we found that the extent of curling (quantified by the curl strain 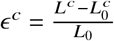, with *L*^*c*^ the curvilinear length of the curl) increased as tissue stress and strain decreased (Fig 6C & D, Movie 11). Remarkably, the curl strain is of the same order of magnitude as in-plane strain, indicative of a strong coupling between in-plane and out-of-plane tissue shapes. These shape changes were in quantitative agreement with predictions from our model parametrized with the mean measured values of the mechanical parameters (*B*, *E*, and *C*_0_) (Fig S5B). In the model, this coupling results from the high spontaneous curvature of the tissue, since decreasing *C*_0_ by an order of magnitude led to a lower change of curl length (Fig S5C). Therefore, we conclude that the coupling between in-plane and out-of-plane monolayer deformation originates from the high spontaneous curvature of the tissue controlled by the asymmetric distribution of myosin II.

**Fig. 6.**
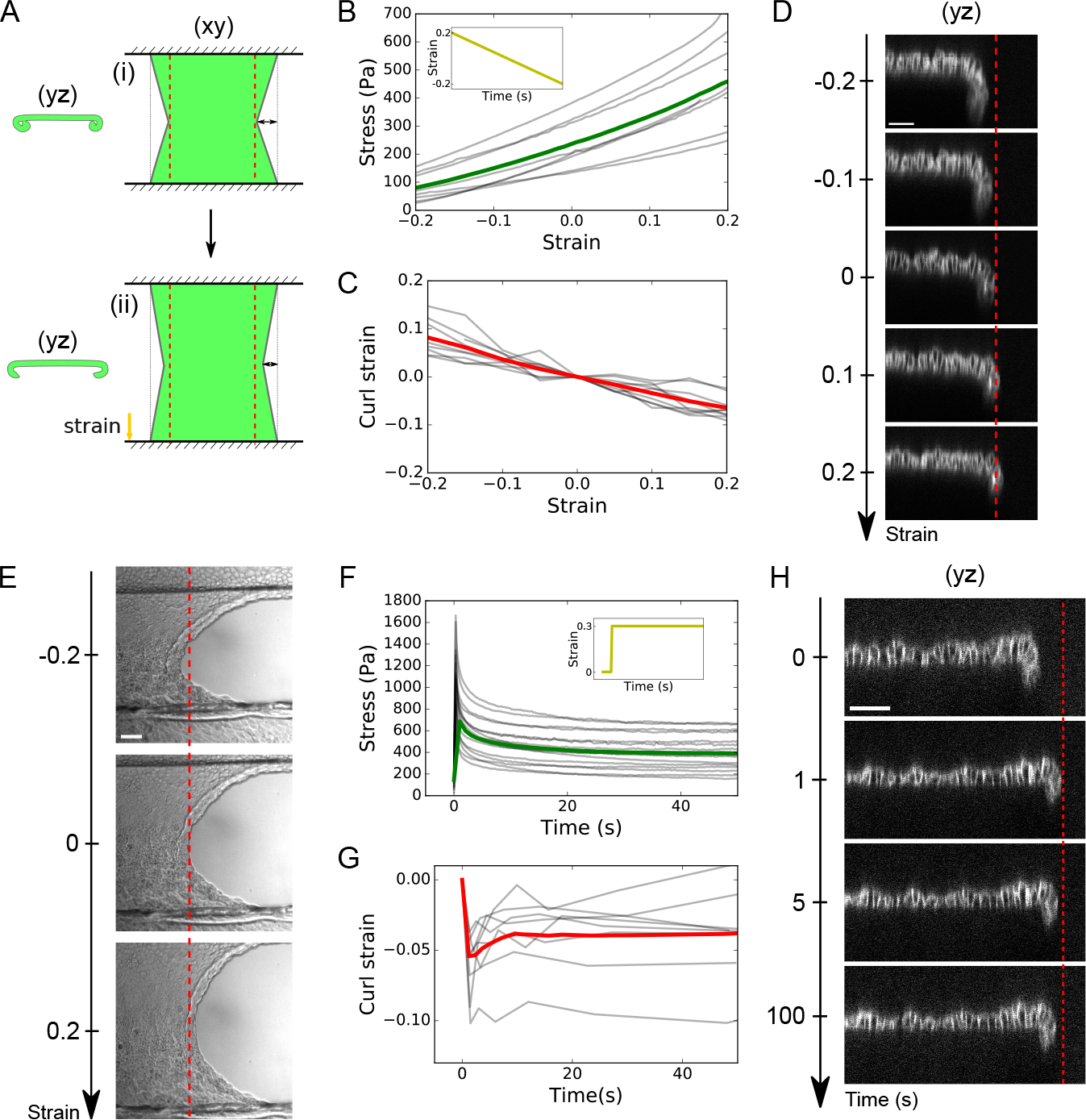
In plane stress and spontaneous curvature interplay to control the extent of curling. (A) Diagram describing the model of 2D elastic sheet used to predict the coupling between in-plane (stretch) and out-of-plane (curling) tissue shape (see main text and Appendix 2). (i) Curled and retracted state naturally reached by the sheet due to spontaneous curvature. For simplicity, the free edge is modeled as a straight diagonal line. (ii) Curled state reached after uni-axial stretching E along the x-axis. (B) Evolution of monolayer stress with in-plane strain during a ramp of compression performed at a rate of 0.5%.s^−1^ after an initial 25% stretch and 5 min period of rest (inset). The grey lines represent 8 individual experiments, the green line represents the average. (C) Evolution of curl strain as a function of in-plane monolayer strain E for the same ramp of deformation as in B. The grey lines represent 9 individual experiments, the red line represents the average. Curl strain is defined in the main text. (D) Typical evolution of MDCK monolayer profile during a ramp of inplane strain. Cell-cell junctions are shown by E-Cadherin-GFP staining. The red dashed line indicates tissue edge at null strain to emphasize the unfurling of the monolayer at positive strain (stretching) and increased curling at negative strain (compression). Scale bar: 50μm. (E) Typical DIC images of monolayer free edge in various strain conditions (−20%, 0% and +20% strain). Scale bar: 50μm. (F) Time evolution of monolayer stress after a 30% step of stretch (inset). The grey lines represent 16 individual datasets, the green line represents the average. (G) Variation of curl strain as a function of time after a 30% step of stretch. The grey lines represent 8 individual datasets, the red line represents the average. (H) Typical evolution of MDCK monolayer profile after a step of in-plane strain. Cell-cell junctions are shown by E-Cadherin-GFP staining. The red dashed line indicates tissue edge immediately after stretch to emphasize the curling of the monolayer as stress relaxes. Scale bar: 50μm.

Interestingly, this coupling also implies that stretching a suspended monolayer causes an increase of its projected width, while compression induces a decrease in tissue width (Fig 6E, Movie 12). This behavior is not due to a bulk negative Poisson ratio (i.e cells in the bulk do not change width, Fig S5C & D) but rather to a behavior of the interface only.

### In-plane stress controls out-of-plane tissue shape

In the above experiment, stress and strain evolve in proportion to one another. Yet our model suggests that in-plane stress controls tissue curling. In previous work, we have shown that, following a step deformation of the monolayer, a substantial portion of stress is relaxed within tens of seconds (20) (Fig 6F). We thus applied a fast step of stretch (+30% strain at 400%.s^−1^), maintaining strain constant while monitoring the curl length as in-plane stress decreases. Immediately upon application of stretch, the length of the curl decreased before gradually increasing and finally reaching a plateau (Fig 6G & 6H, Movie 13). The time-scale of this retraction was commensurate with that of in-plane stress relaxation (Fig 6F & 6G). Therefore, the coupling between in-plane (xy) and out-of-plane (xz or yz) tissue shape is controlled by the in-plane tissue stress, rather than the deformation.

## Discussion

Active torques have long been known to induce out-of-plane deformation during epithelial morphogenesis. However, because of a lack of direct measurements, their importance relative to other forces in epithelia has been challenging to assess. Here, we provide a direct quantification of out-of-plane stresses in epithelial monolayers by measuring the stress required to flatten naturally curled M DCK t issues *in vitro*. Curling stresses were lower but on the same order of magnitude as the in-plane pre-tension (≃ 30 Pa to flatten one third of the curled length (Fig 3D), compared with ≃ 250 Pa in-plane pre-tension (11)). In other words, the bending energy and stretching energy – both controlled by myosin molecular motors - are in the same range. Furthermore, direct measurement of the bending stiffness of the tissue revealed it to be low, as expected from thin plate theory. An immediate consequence of these properties is that the spontaneous curvature of the tissue is high, with a radius of curvature on the same order of magnitude as the monolayer thickness. Such high spontaneous curvature is rare in inert materials, although at a different scale it can be observed in the cell membrane under the action of specialized proteins (1).

Our measurements can be used to test the predictions of continuous mechanical models in specific biological contexts, for example to predict the shape of early pancreatic tumour lesions (10). Indeed, depending on the duct radius, tumour lesions evaginate or invaginate and the critical radius *R*\* was proposed to be equal to the ratio between the bending modulus and the bending moment of the epithelium. Using our measurements yields 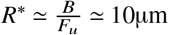, which is on the order of magnitude of the *in vivo* measurements (with *F*_*u*_ ≃ 30*nN* the typical force required to unfurl a commensurate section of epithelium), which suggests that the mechanical properties measured *in vitro* could be relevant to epithelia in general.

The second consequence of high out-of-plane stresses is that they induce a coupling between tissue deformations in-plane and out-of-plane. We show here that changes in boundary conditions in the plane result in a visible change of the curled length at the free edge. In other words, the deformations observed out-of-plane were on the same order of magnitude as those imposed in-plane. Although we cannot exclude other potential mechanical feedback (for example myosin flows in response to the variation of in-plane stresses), this effect is likely independent of cell signaling. Indeed, curling decreased immediately after the tissue was stretched at very high strain rate leading to deformation on a sub-second timescale (Fig 6H). Furthermore, our mechanical model could predict changes of tissue shape out-of-plane (curled length) in response to a ramp of in-plane deformation by minimizing the elastic energy of a 2D sheet endowed with constant and isotropic spontaneous curvature. This indicates that the coupling exists without any change of the tissue spontaneous curvature, which could occur in response to signaling. Such mechanical coupling could have implications *in vivo* as it was shown recently that disruption of in-plane tension anisotropy in the Drosophila embryo can prevent mesoderm invagination driven by apical constriction (30), while compression generated by cell migration is required to correctly shape the optic cup alongside basal recruitment of Myosin II in zebrafish (31). Since in-plane stresses can be transferred through long-range mechanical interactions (32, 33), our measurements further emphasize a potential role for tissue-scale force generation in the shaping of epithelia.

Importantly, rather than being a curiosity restricted to *in vitro* settings, we have shown that curling also occurs *in vivo* as part of developmental morphogenesis, during *Drosophila* leg eversion. In this process, the peripodial epithelium detaches from its matrix, fractures, and curls as it retracts to allow leg extension. Since the apical side of the peripodial epithelium faces the apical surface of the leg columnar epithelium, basal curling of the peripodial epithelium may facilitate leg eversion by limiting frictional forces between the two tissues. Furthermore, epithelial curling has also been reported in pathological conditions. During age-related macular degeneration, the retinal epithelium detaches from its basement membrane, ruptures and retracts, leading to partial blindness of the patient (34). The curling that occurs in this case might have a negative impact by amplifying the opening of the retina (35). Thus, we expect epithelial curling to be involved in other physiological or pathological phenomena where a substrate-free epithelial monolayer exhibits a free edge and myosin II polarization creates a spontaneous curvature of the tissue.

## Materials and methods

### MDCK cell culture

MDCK cells were cultured at 37°C in an atmosphere of 5% *CO*_2_. Cells were passaged at a 1:5 ratio every 3-4 days using standard cell culture protocols and disposed of after 25 passages. The culture medium was composed of high glucose DMEM (Thermo Fisher Scientific) supplemented with 10% FBS (Sigma) and 1% penicillin-streptomycin (Thermo Fisher Scientific).

### Generation of suspended MDCK tissues

Suspended monolayers of MDCK cells were generated as described in (15, 23). Briefly, a drop of collagen was placed between two test rods and left to dry at 37°C to form a solid scaffold. This collagen was then rehydrated before cells were seeded onto it and cultured for 72 hours. Immediately before each experiment, the collagen scaffold was removed via enzymatic digestion with collagenase leaving a monolayer without extra-cellular matrix suspended in between the two test rods.

## Supplementary materials and methods

Additional material and methods can be found in the Supplementary Information text.

## ACKNOWLEDGEMENTS

The authors wish to acknowledge present and past members of the Charras, Kabla and Suzanne labs for stimulating discussions. They also thank Emmanuel Martin, Richard Thorogate, Duncan Farquharson and Simon Townsend for their precious help in designing experiments. J.F. and P.R. were funded by BBSRC grant (BB/M003280 and BB/M002578) to G.C. and A.K. J.F was funded by an EMBO Short-term fellowship (number 7824) for the work carried out on peripodial epithelium. A.P was funded by a consolidator grant to M.S from the European Research Council (EPAF, agreement 648001). A.L. was supported by an EMBO long term post-doctoral fellowship (number 29-2016). J.F, T.W., N.G, A.L., N.K., and G.C. were supported by a consolidator grant from the European Research Council to G.C (MolCellTissMech, agreement 647186).

## References

1. Joshua Zimmerberg and Michael M Kozlov. How proteins produce cellular membrane curvature. Nature reviews Molecular cell biology, 7(1):9, 2006.

2. Utpal Nath, Brian CW Crawford, Rosemary Carpenter, and Enrico Coen. Genetic control of surface curvature. Science, 299(5611):1404–1407, 2003.

3. Kristin Sherrard, François Robin, Patrick Lemaire, and Edwin Munro. Sequential activation of apical and basolateral contractility drives ascidian endoderm invagination. Current Biology, 20(17):1499–1510, 2010.

4. Bruno Monier, Melanie Gettings, Guillaume Gay, Thomas Mangeat, Sonia Schott, Ana Guarner, and Magali Suzanne. Apico-basal forces exerted by apoptotic cells drive epithelium folding. Nature, 518(7538):245, 2015.

5. Thomas Lecuit and Pierre-Francois Lenne. Cell surface mechanics and the control of cell shape, tissue patterns and morphogenesis. Nature reviews Molecular cell biology, 8(8):633, 2007.

6. Esther J Pearl, Jingjing Li, and Jeremy BA Green. Cellular systems for epithelial invagination. Philosophical Transactions of the Royal Society B: Biological Sciences, 372(1720): 20150526, 2017.

7. Verena Kölsch, Thomas Seher, Gregorio J Fernandez-Ballester, Luis Serrano, and Maria Leptin. Control of drosophila gastrulation by apical localization of adherens junctions and rhogef2. Science, 315(5810):384–386, 2007.

8. Adam C Martin, Matthias Kaschube, and Eric F Wieschaus. Pulsed contractions of an actin–myosin network drive apical constriction. Nature, 457(7228):495, 2009.

9. Mahim Misra, Basile Audoly, Ioannis G Kevrekidis, and Stanislav Y Shvartsman. Shape transformations of epithelial shells. Biophysical journal, 110(7):1670–1678, 2016.

10. Hendrik A Messal, Silvanus Alt, Rute MM Ferreira, Christopher Gribben, Victoria Min-Yi Wang, Corina G Cotoi, Guillaume Salbreux, and Axel Behrens. Tissue curvature and apicobasal mechanical tension imbalance instruct cancer morphogenesis. Nature, 566(7742): 126, 2019.

11. Tom PJ Wyatt, Jonathan Fouchard, Ana Lisica, Nargess Khalilgharibi, Buzz Baum, Pierre Recho, Alexandre J Kabla, and Guillaume T Charras. Actomyosin controls planarity and folding of epithelia in response to compression. Nature materials, pages 1–9, 2019.

12. Edouard Hannezo, Jacques Prost, and Jean-Francois Joanny. Theory of epithelial sheet morphology in three dimensions. Proceedings of the National Academy of Sciences, 111 (1):27–32, 2014.

13. Stephanie Höhn, Aurelia R Honerkamp-Smith, Pierre A Haas, Philipp Khuc Trong, and Raymond E Goldstein. Dynamics of a volvox embryo turning itself inside out. Physical review letters, 114(17):178101, 2015.

14. Guillaume Salbreux and Frank Jülicher. Mechanics of active surfaces. Physical Review E, 96(3):032404, 2017.

15. Andrew R Harris, Loic Peter, Julien Bellis, Buzz Baum, Alexandre J Kabla, and Guillaume T Charras. Characterizing the mechanics of cultured cell monolayers. Proceedings of the National Academy of Sciences, 109(41):16449–16454, 2012.

16. Sri Ram Krishna Vedula, Hiroaki Hirata, Mui Hoon Nai, Yusuke Toyama, Xavier Trepat, Chwee Teck Lim, Benoit Ladoux, et al. Epithelial bridges maintain tissue integrity during collective cell migration. Nature materials, 13(1):87, 2014.

17. Laura Casares, Romaric Vincent, Dobryna Zalvidea, Noelia Campillo, Daniel Navajas, Marino Arroyo, and Xavier Trepat. Hydraulic fracture during epithelial stretching. Nature materials, 14(3):343, 2015.

18. Ernest Latorre, Sohan Kale, Laura Casares, Manuel Gómez-González, Marina Uroz, Léo Valon, Roshna V Nair, Elena Garreta, Nuria Montserrat, Aránzazu del Campo, et al. Active superelasticity in three-dimensional epithelia of controlled shape. Nature, 563(7730):203, 2018.

19. Medhavi Vishwakarma, Jacopo Di Russo, Dimitri Probst, Ulrich S Schwarz, Tamal Das, and Joachim P Spatz. Mechanical interactions among followers determine the emergence of leaders in migrating epithelial cell collectives. Nature communications, 9(1):3469, 2018.

20. Nargess Khalilgharibi, Jonathan Fouchard, Nina Asadipour, Ricardo Barrientos, Maria Duda, Alessandra Bonfanti, Amina Yonis, Andrew Harris, Payman Mosaffa, Yasuyuki Fujita, et al. Stress relaxation in epithelial monolayers is controlled by the actomyosin cortex. Nature Physics, page 1, 2019.

21. Manouk Abkarian, Gladys Massiera, Laurence Berry, Magali Roques, and Catherine Braun-Breton. A novel mechanism for egress of malarial parasites from red blood cells. Blood, 117(15):4118–4124, 2011.

22. Stéphane Douezan, Matthieu Wyart, Françoise Brochard-Wyart, and Damien Cuvelier. Curling instability induced by swelling. Soft Matter, 7(4):1506–1511, 2011.

23. Andrew R Harris, Julien Bellis, Nargess Khalilgharibi, Tom Wyatt, Buzz Baum, Alexandre J Kabla, and Guillaume T Charras. Generating suspended cell monolayers for mechanobiological studies. Nature protocols, 8(12):2516, 2013.

24. Matthew C Gibson and Gerold Schubiger. Drosophila peripodial cells, more than meets the eye? Bioessays, 23(8):691–697, 2001.

25. Martin J Milner, Alison J Bleasby, and Susan L Kelly. The role of the peripodial membrane of leg and wing imaginal discs ofdrosophila melanogaster during evagination and differentiation in vitro. Wilhelm Roux’s archives of developmental biology, 193(3):180–186, 1984.

26. Silvia Aldaz, Luis M Escudero, and Matthew Freeman. Dual role of myosin ii during drosophila imaginal disc metamorphosis. Nature communications, 4:1761, 2013.

27. Amsha Proag, Bruno Monier, and Magali Suzanne. Physical and functional cell-matrix uncoupling in a developing tissue under tension. Development, 146(11):dev172577, 2019.

28. Anastasiya Trushko, Ilaria Di Meglio, Aziza Merzouki, Carles Blanch-Mercader, Shada Abuhattum, Jochen Guck, Kevin Alessandri, Pierre Nassoy, Karsten Kruse, Bastien Chopard, et al. Buckling of epithelium growing under spherical confinement. bioRxiv, page 513119, 2019.

29. Mathieu Poujade, Erwan Grasland-Mongrain, A Hertzog, J Jouanneau, Philippe Chavrier, Benoît Ladoux, Axel Buguin, and Pascal Silberzan. Collective migration of an epithelial monolayer in response to a model wound. Proceedings of the National Academy of Sciences, 104(41):15988–15993, 2007.

30. Soline Chanet, Callie J Miller, Eeshit Dhaval Vaishnav, Bard Ermentrout, Lance A Davidson, and Adam C Martin. Actomyosin meshwork mechanosensing enables tissue shape to orient cell force. Nature communications, 8:15014, 2017.

31. Jaydeep Sidhaye and Caren Norden. Concerted action of neuroepithelial basal shrinkage and active epithelial migration ensures efficient optic cup morphogenesis. Elife, 6:e22689, 2017.

32. Kapil Bambardekar, Raphaël Clément, Olivier Blanc, Claire Chardès, and Pierre-François Lenne. Direct laser manipulation reveals the mechanics of cell contacts in vivo. Proceedings of the National Academy of Sciences, 112(5):1416–1421, 2015.

33. Mahamar Dicko, Pierre Saramito, Guy B Blanchard, Claire M Lye, Bénédicte Sanson, and Jocelyn Étienne. Geometry can provide long-range mechanical guidance for embryogenesis. PLoS computational biology, 13(3):e1005443, 2017.

34. Rama D Jager, William F Mieler, and Joan W Miller. Age-related macular degeneration. New England Journal of Medicine, 358(24):2606–2617, 2008.

35. Neil M Bressler, Susan B Bressler, and Stuart L Fine. Age-related macular degeneration. Survey of ophthalmology, 32(6):375–413, 1988.

