## Supplementary information for "Curling of epithelial monolayers reveals coupling between active bending and tissue tension"

### Supplementary information for
### Curling of epithelial monolayers reveals coupling between active bending and tissue tension

**Jonathan Fouchard, Tom P.J. Wyatt, Amsha Proag, Ana Lisica, Nargess Khalilgharibi, Pierre Recho, Magali Suzanne, Alexandre Kabla, and Guillaume Charras**

#### Supplementary methods

**Fly stock.** Flies were grown under standard culture techniques. A Sqh-TagRFPt[9B] knock-in line was used for imaging of Myosin II homolog in *Drosophila* during peripodial epithelium retraction (1). A Flytrap line Vkg-GFP[G0454] was used for imaging the dynamics of the collagen (extra-cellular matrix) during leg eversion (2). The Armadillo-GFP line is from Bloomington (number 8556).

**Leg disc preparation.** Leg discs were dissected from white pupae + 2 hours after puparium formation in Schneider's insect medium (Sigma-Aldrich) supplemented with 15% fetal calf serum, 0.5% penicillin-streptomycin and 2 µg/mL 20-hydroxyecdysone (Sigma-Aldrich, H5142). Leg discs were transferred onto a glass slide in 13.5 µL of this medium and confined between a 120 µm-deep double-sided adhesive spacer (Secure-SealTM from Sigma-Aldrich) and a glass coverslip placed on top of the spacer. Halocarbon oil was added to the sides of the spacer to prevent evaporation. To visualize cell shapes during peripodial epithelium retraction, cell membranes were labeled after dissection and before imaging via a 10-minute incubation with Far red CellMask plasma membrane stain according to manufacturer instructions (Thermo Fisher Scientific). To prevent collagen remains from interfering with peripodial epithelium curling, a collagenase treatment (0.18 units/mL) was applied for 10 minutes simultaneously with CellMask incubation.

**Confocal imaging of peripodial epithelium.** Retraction and curling of peripodial epithelia were imaged at 24°C on an inverted confocal laser scanning microscope (LSM-880, Zeiss) equipped with an Airyscan detector and a 40X objective (C-Apochromat, NA=1.2, Zeiss). Images were acquired at a rate of one z-stack every 2-5min and z-slices were spaced by 0.5µm. Airyscan images were then reconstructed using the Airyscan processing module of the Zen Black software (Zeiss). Movies of the retraction were then generated using Imaris (Bitplane).

**MDCK cell lines.** MDCK-E-Cadherin-GFP cell lines (generated as described in (3)) were cultured in presence of 250ng/ml puromycin in the culture medium. MDCK NMHCIIA-GFP and MDCK NMHCIIB-GFP were generated as described in (4). Cells were then cultured in presence of G418 (1mg/ml) in the culture medium.

**Imaging and quantification of Myosin II-GFP distribution in MDCK epithelia and Sqh-RFP distribution in *Drosophila* peripodial epithelia.** To quantify the anisotropy of Myosin II along the apico-basal axis of suspended MDCK cells, z-stacks of MDCK monolayers expressing NMHCIIA–GFP or NMHCIIB–GFP were acquired using a high numerical aperture silicon oil 40X objective (UPLSAPO S, NA=1.25, Olympus) mounted on an Olympus IX83 inverted microscope equipped with a scanning laser confocal head (Olympus FV1200). Apical, medial and basal surfaces were segmented from images of the plasma membrane stained with Far red CellMask (Thermo Fisher Scientific). A z-projection of 2 focal planes separated by 1µm was generated at each position (apical, medial, or basal) to compensate for the slight tilt of the cells along the z-axis. Myosin II average intensity was then measured within the segmented regions. To account for background fluorescence, the average intensity of a focal plane above the apical zone was subtracted from the measurement. We also noticed the existence of a gradient of intensity along the thickness axis in the images of the plasma membrane stain. Such gradient was consistent with a decay of the incident and emitted light in the deeper regions of the cells. We thus took this effect into account in our calculation of the average intensity in the basal compartment as follows: $I_c^{bas} = (I^{bas} - I^n)\frac{I_m^{ap}}{I_m^{bas}}$ and $I_c^{ap} = I^{ap} - I^n$, where $I_c^{ap}$ and $I_c^{bas}$ are the corrected intensities of apical and basal Myosin II respectively, $I^n$ is the intensity of the background (noise), $I^{ap}$ and $I^{bas}$ are the measured Myosin II intensities in the apical and basal domains respectively, and $I_m^{ap}$ and $I_m^{bas}$ are the intensities of the CellMask membrane marker in the apical and basal domains respectively.

**Measurement of tissue curvature.** The curvature of MDCK monolayers or *Drosophila* peripodial epithelia along their contour length was determined as follows. First, the coordinates of the midpoint of each baso-lateral junction was manually determined from a profile view of the epithelium. A spline (*interpolate.splrep* of the *Scipy* Python library) was then fitted to these points to obtain the full monolayer profile. The local curvature $\kappa$ along the monolayer contour length was subsequently computed every 1µ$m$ from the interpolating spline given parametrically (y(t), z(t)): $\kappa = \frac{|y''z' - y'z''|}{(z'^2 + y'^2)^{\frac{3}{2}}}$. The spontaneous curvature of the tissue was defined as the average curvature within the tip region of the tissue (green region on Fig S1B). Note that the definition of monolayer tip is dependent on monolayer shape. Indeed, two categories of shapes could be identified based on tissue curvature. In some cases, the monolayer was only bent at the very edge (e.g. for Y27632 treated tissues), whereas in other cases the monolayer curled up on itself (see Fig 1F). In the first case, the curvature was defined within the region after the last maxima of $z(l)$, with z the z-coordinate of the profile and $l$ the curvilinear distance along the monolayer contour. In the latter case, the curvature was computed within the region where the y-coordinate of the tissue profile $y(l)$ evolved non-monotonously with $l$.

**Mechanical manipulation of MDCK epithelial tissues for bending forces measurements.** Local unfolding of MDCK curled monolayers were performed as follows. A glass capillary was pulled with a micropipette puller (Narishige) and its tip was cut and glued to a second stiff glass capillary which was previously bent to accommodate the geometry of the setup. The stiffness $k$ of the device was calibrated by pressing it against a Nitinol wire of known stiffness (see (3)) in order to serve as a force cantilever. Before the experiment, the force cantilever was approached in the vicinity of the curled tissue free edge of the MDCK monolayer. To unfold the tissue, a ramp of displacement $D(t)$ was imposed at the base of the device at a rate of 0.5 $\mu m.s^{-1}$ through a motorized platform (M-126.DG1 controlled through a C-863 controller, Physik Instrumente, via a Labview program). The displacement of the tip of the cantilever $\Delta(t)$ was then measured from confocal images after segmentation through the *Triangle* thresholding algorithm available in Fiji (5). The force required to unfurl the MDCK monolayer was then equal to: $F(t) = k.(D(t) - \Delta(t)) = k.\delta(t)$ with $\delta(t)$ the deflection of the cantilever.

**Mechanical manipulation of MDCK epithelial tissues in the tissue plane: stretching and compression.** Tissue-scale mechanical deformations in the plane of MDCK monolayers along the x-axis were applied as described in (6). Briefly, a custom-made adaptor was wedged in the top end of the hinged arm of the stretching device. The adaptor was connected to a 2-D manual micromanipulator mounted on a motorized platform (M-126.DG1 controlled through a C-863 controller, Physik Instrumente). Then, the tissues were deformed by moving the motorized platform via a custom-made Labview program (National Instruments). To image the same yz-profile of the tissue during stretching, that portion of the tissue needs to stay immobile in the microscope reference frame. For that, the motorized platform was interfaced with the microscope stage (PS3J100, Prior Scientific Instruments) through a custom-made Labview program as follows. If the tissue of length $L_0$ (i.e the initial distance between coverslips) was stretched by a length $l$ and a profile located at a distance $x < L_0$ from the static coverslip was imaged, the stage was then moved by $-\frac{lx}{L_0}$ to compensate for the change in length of the tissue. The profile chosen in our experiments was the one where the deflection of the tissue was maximum, corresponding to $x \simeq \frac{L_0}{2}$.

**Confocal imaging of MDCK epithelial tissues during mechanical manipulation.** Tissues were imaged at 37°C in a humidified atmosphere with 5% CO2. The imaging medium consisted of DMEM without phenol red supplemented with 10% FBS. Profile views of the curled tissues during mechanical manipulation were obtained using a high numerical aperture silicon oil 30X objective (UPLSAPO S, NA=1.05, Olympus) mounted on an Olympus IX83 inverted microscope equipped with a scanning laser confocal head (Olympus FV1200). Each image consisted in roughly 100 slices spaced by $1\mu m$. Time series were acquired with an interval of ≃1s. To visualize the tip of the cantilever, AlexaFluor-647-conjugated dextran (10,000 MW, Thermo Fisher Scientific, added at $20\mu g.mL^{-1}$) was added to the medium. In addition to staining the medium, we noticed that dextran accumulated at the cantilever tip, allowing for easy segmentation. While we do not know the cause of this accumulation, we hypothesize that it is due to preferential binding to the oxidized region of the pulled capillary.

**Laser ablation of MDCK epithelial tissues.** Laser ablation and subsequent imaging of MDCK tissues was carried out with a LSM880 scanning laser confocal system mounted on an inverted Axio Observer microscope stand (Zeiss). Ablation was performed through an infrared Ti:Sapphire laser tuned at 980nm (200 mW emission) over the thickness of the sample in rectangular regions (150 x 15 $\mu m^2$, 16 x 2 cells) situated in the centre of suspended MDCK monolayers. Confocal stacks were acquired through a 40X objective by exciting the fluorophores at the appropriate wavelength (LD C-Apochromat, NA=1.1, Zeiss). Each stack consisted in roughly 50 slices spaced by $1\mu m$. Time series were acquired every 20-60 seconds for 12 minutes minimum.

**Statistics and data analysis.** All data and statistical analysis were performed using the Python language environment and its scientific libraries (NumPy (7), SciPy (8)). Graphs were plotted using the Python library Matplotlib (9). Basic image processing was carried out with the Fiji package (5). The statistical significance between the different samples was assessed using the Mann-Whitney U test. Significance symbols were defined as follows: (****, $p<0.0001$), (***, $p<0.001$), (**, $p<0.01$), (*, $p<0.05$). All boxplots show the mean value (square), the median value (central bar), the first and third quartile (bounding box) and the range (whiskers) of the distribution. All code is available from the corresponding author upon reasonable request.

#### Appendix 1 : Method to estimate the bending modulus of epithelial monolayers

We have designed a method to measure the force necessary to unfurl a curled suspended MDCK epithelial monolayer (Fig 3). Here, we describe how we could calculate the 2-dimensional bending modulus $B$ of such tissue from these measurements, using an energetic method.

As the force cantilever is displaced along the y-axis perpendicular to tissue free edge, it 1) unfurls the previously curled tissue and 2) stretch part of the flat tissue in the bulk. The work $W_f$ performed by the cantilever along its displacement is then equal to the sum of the mechanical energies deforming the monolayer: $W_c$ the bending energy necessary to change the curvature of the curl from $C_0$ to $C_1$, and $W_s$ the stretching energy of the area in the area in the bulk deflected of $\delta_b$.

$$W_f = W_c + W_s \qquad [1]$$

- The work of the cantilever corresponding to a course $\Delta_c$ of its tip ($\Delta_c = D - \delta$ in the main manuscript) and the respective applied force $F_c$ is :

$$W_f = \frac{1}{2} F_c \Delta_c \qquad [2]$$

- The bending energy stored in the curl is:

$$W_c = \frac{1}{2} B L^c w_c (C_1^2 - C_0^2) \qquad [3]$$

where $B$ is the bending modulus, $L^c$ the length of the curl and $w_c$ the width of the deflected tissue.

- The stretching energy of the bulk is :

$$W_s = \frac{1}{2} E w_c L_b (\frac{\delta_b}{L_b}^2) \simeq \frac{1}{2} E \delta_b{}^2 \qquad [4]$$

where $E$ is the 2-D elastic modulus of the tissue $E = E_{3D}.h$, $h$ the monolayer thickness and $L_b$ the length over which the bulk tissue is deflected.

The measurements of $L^c$, $w_c$, $h$, $L_b$ and $\delta_b$ were performed manually from confocal microscopy images. $\Delta_c$ was measured through automatic segmentation of confocal images (see Methods). The value of $E_{3D}$ used was obtained from force-extension tests (Fig 6B) (6). $C_0$ and $C_1$ were computed as the average curvatures in the curled region before and after application of force (Fig 3F).

#### Appendix 2 : Model of 2D elastic thin sheet to capture the coupling between in-plane and out-of-plane epithelial monolayer shape

**Summary.** We model the epithelial monolayer as an elastic rectangular bi-dimensional sheet clamped to two infinitely rigid plates. One of the plates can be displaced uni-axially in the direction parallel to the free edge of the sheet, while the other one is fixed. The clamped sides of the sheet are of length $w$ whereas the free edges of the sheet are of length $L_0$. The sheet has a 2D bending modulus $B$, a 2-D stiffness $E$ and is endowed with a spontaneous curvature $C_0$.

The ultimate shape of the tissue will be determined by a balance between curling - which relax the spontaneous curvature while deflecting the tissue inwards - and stretching of the free edge which varies according to the position of the plates and the amount of curled tissue (see main manuscript). Our aim is to determine the maximum deflection of the tissue $d$ caused by curling at the tissue free edge according to monolayer boundary conditions.

**Calculation of the deflection of the sheet.** We start from a hypothetical initial configuration where the sheet is fully flattened (Fig S6,(i)). The energy associated to this state corresponds to the energy required to cancel the spontaneous curvature $C_0$:

$$E_0 = \frac{1}{2} w L_0 B C_0^2 \qquad [5]$$

We then assume the sheet to be free to curl, which will deflect the tissue inwards by a distance $d$ (Fig S6, (ii)). Yet, due to the clamped sides of the sheet, the free edge of the tissue gets stretched. We approximate the free edge interface by two straight segments. (This is a simplification since the free edge of the monolayer possess an arc shape in our experiments.) The curled region of area $\frac{1}{2} d L_0$ has now a curvature $C_0$ (second term in the equation below). We assume that this area is then stretched with a strain $\epsilon_c(d, L_0)$ corresponding to the extension of the free edge from a length $L_0$ to a length $2l$ (third term in the equation below, Fig S6, (ii)). The fourth term in the equation below represents the stretching energy of the bulk (of strain $\epsilon$). However, this term will not play a role in the final definition of the shape of the free edge. The energy $E_1$ of the sheet can then be written as :

$$E_1(\epsilon, d) = E_0 - \frac{1}{2} d L_0 B C_0^2 + \frac{1}{2} d L_0 E \epsilon_c^2 + \frac{1}{2} w L_0 E \epsilon^2 \qquad [6]$$

We can rewrite $\epsilon_c$ as a function of $l$ (where $2l$ is the new length of the free edge) and $L_0$ :

$$\epsilon_c = \epsilon + \frac{2l}{L_0} - 1 = \epsilon + \sqrt{4(\frac{d}{L_0})^2 + 1} - 1 \qquad [7]$$

We then search for $d$ which minimizes $E_1$ (Fig S6, (iii)) :

$$\frac{\partial E_1(\epsilon,d)}{\partial d}|_\epsilon = 0 \quad [8]$$

This leads to :

$$\frac{BC_0^2}{E} = (\sqrt{4(\frac{d}{L_0})^2+1}+\epsilon-1)^2 + 8(\frac{d}{L_0})^2\frac{\sqrt{4(\frac{d}{L_0})^2+1}+\epsilon-1}{\sqrt{4(\frac{d}{L_0})^2+1}} \quad [9]$$

We know from experiments that $\frac{d}{L_0} \simeq \frac{1}{2}$, so we can develop $4(\frac{d}{L_0})^2$ around 1, which leads to:

$$\sqrt{4(\frac{d}{L_0})^2+1} \simeq -\frac{1}{\sqrt{2}}(\frac{d}{L_0})^4 + \frac{5}{2\sqrt{2}}(\frac{d}{L_0})^2 + 1 \quad [10]$$

Removing the highest order in the development of Equation 9, we obtain :

$$\frac{BC_0^2}{E} = 8(\frac{d}{L_0})^2\epsilon + 17(\frac{d}{L_0})^4 \quad [11]$$

Equation 11 therefore predicts the evolution of monolayer deflection $d$ (corresponding to the curled tissue length) as a function of sample initial length $L_0$ and the deformation $\epsilon$ through only three parameters: the 2-D stiffness $E$, the bending modulus $B$ and the spontaneous curvature $C_0$. Importantly, all these parameters could be measured in our setup.

According to Equation 11, on the first order, for $\epsilon = 0$, $d$ should be proportional to $L_0$ : $d = \alpha L_0$, which we could observe experimentally with the fitting pre-factor $\alpha_{exp} = 0.50$ (Fig 5F), close to the predicted pre-factor $\alpha_{th} = 0.78$.

Equation 11 also predicts how $d$ evolves when the boundary conditions of the tissue (i.e $\epsilon$) are changed :

$$\frac{d}{L_0} = \sqrt{\frac{\sqrt{64\epsilon^2 + 68\frac{BC_0^2}{E}} - 8\epsilon}{34}} \quad [12]$$

Figure S5B shows the evolution of the deflection $d$ as a function of in-plane tissue strain $\epsilon$. The model prediction is in accordance with the variations of deflection measured without any fitting parameter.

#### Supplementary references


1. A Proag, B Monier, M Suzanne, Physical and functional cell-matrix uncoupling in a developing tissue under tension. *Development* **146**, dev172577 (2019).
2. X Morin, R Daneman, M Zavortink, W Chia, A protein trap strategy to detect gfp-tagged proteins expressed from their endogenous loci in drosophila. *Proc. Natl. Acad. Sci.* **98**, 15050–15055 (2001).
3. AR Harris, et al., Characterizing the mechanics of cultured cell monolayers. *Proc. Natl. Acad. Sci.* **109**, 16449–16454 (2012).
4. N Khalilgharibi, et al., Stress relaxation in epithelial monolayers is controlled by the actomyosin cortex. *Nat. Phys.*, 1 (2019).
5. J Schindelin, et al., Fiji: an open-source platform for biological-image analysis. *Nat. methods* **9**, 676 (2012).
6. TP Wyatt, et al., Actomyosin controls planarity and folding of epithelia in response to compression. *Nat. materials*, 1–9 (2019).
7. S Van Der Walt, SC Colbert, G Varoquaux, The numpy array: a structure for efficient numerical computation. *Comput. Sci. & Eng.* **13**, 22 (2011).
8. E Jones, T Oliphant, P Peterson, , et al., Scipy: Open source scientific tools for python. (2001).
9. JD Hunter, Matplotlib: A 2d graphics environment. *Comput. science & engineering* **9**, 90 (2007).

### Figure S1

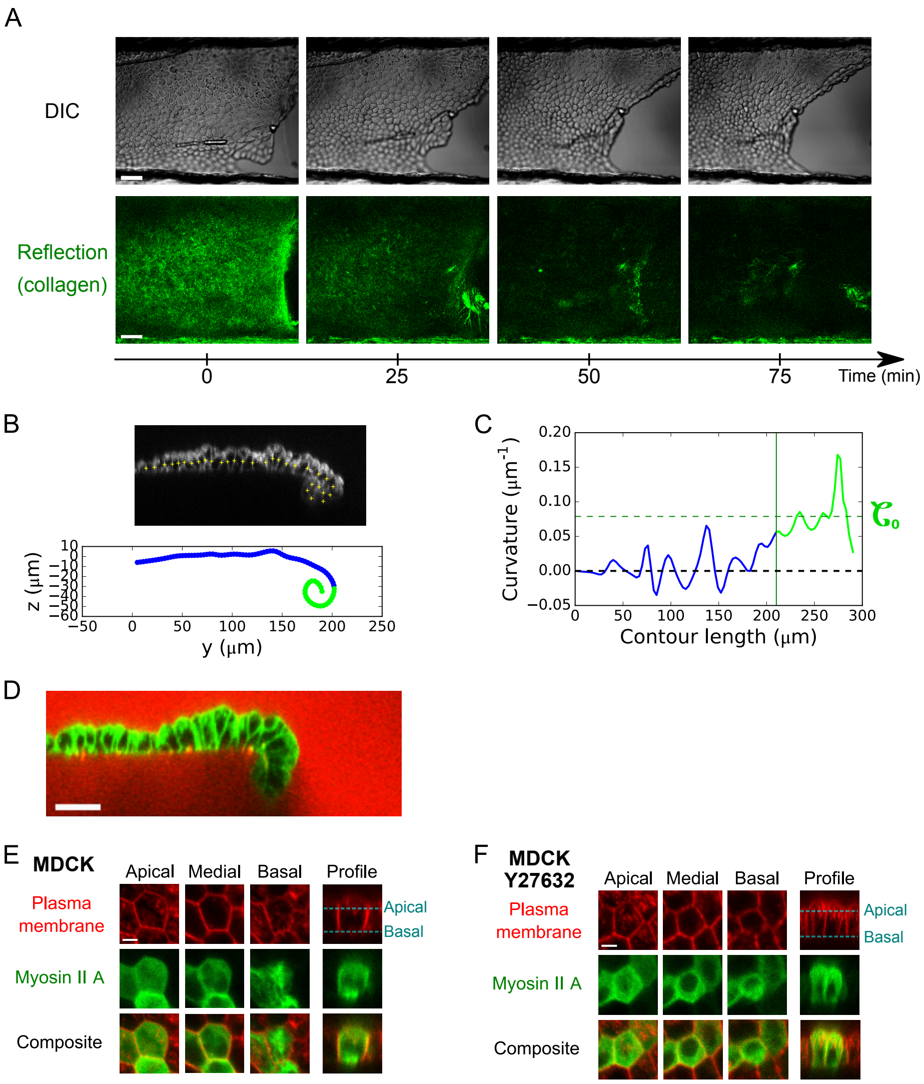


**Fig.S 1.** (A) Time-series showing the shape changes of the free edge of an MDCK monolayer during the digestion of the collagen substrate with collagenase (N=14). Top: DIC imaging shows changes of tissue shape. Bottom: Confocal reflection microscopy imaging shows the dynamics of collagen digestion. Scale bars: 50μm. (B) Extraction of the monolayer profile contour. Top: Profile view of an MDCK monolayer obtained through confocal scanning laser imaging. Cell membranes are marked with CellMask. The coordinates of the midpoint of each baso-lateral junction are manually positioned. Bottom: A spline is fitted to these coordinates to define the monolayer contour. (C) The local curvature is calculated along the monolayer contour length in 1μm steps (yellow crosses). The spontaneous curvature of the tissue is determined as the average curvature within the tip region of the tissue free edge (green part in B and C, see Methods). In this example, the tip is defined as the region after the y-coordinate of the tissue profile y($l$), where $l$ is the contour length of the tissue, evolved non-monotonously with $l$ (see green region in B, bottom). (D) Scanning laser confocal image showing the profile view (yz-plane) of a representative suspended MDCK monolayer. Cell membranes are marked with CellMask (green). The medium is marked with Dextran-Alexa647 (red). Note that no medium can be observed inside the curl, indicative of a very high curvature of the monolayer. (E) Localisation of Myosin II A-GFP in the apical, medial and basal regions of a substrate-free MDCK monolayer. Plasma membrane is marked with CellMask. Scale bar: 5μm. (F) Localisation of Myosin II A-GFP in the apical, medial and basal regions of a substrate-free MDCK monolayer treated with 25μM Y27632. Plasma membrane is marked with CellMask. Scale bar: 5μm.

#### Figure S2

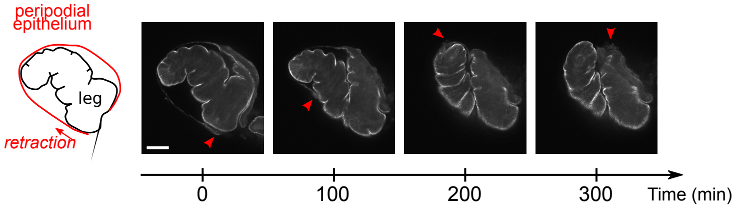


**Fig.S 2.** Low magnification confocal imaging of a *Drosophila* leg undergoing eversion. The outer peripodial epithelium (in red on the diagram) retracts, while the leg epithelium (center) changes shape. Red arrowheads indicate the retracting peripodial epithelium. Cell junctions are marked with Armadillo-GFP (*Drosophila* homolog of β-catenin). Scale bar: 50μm.

#### Figure S3

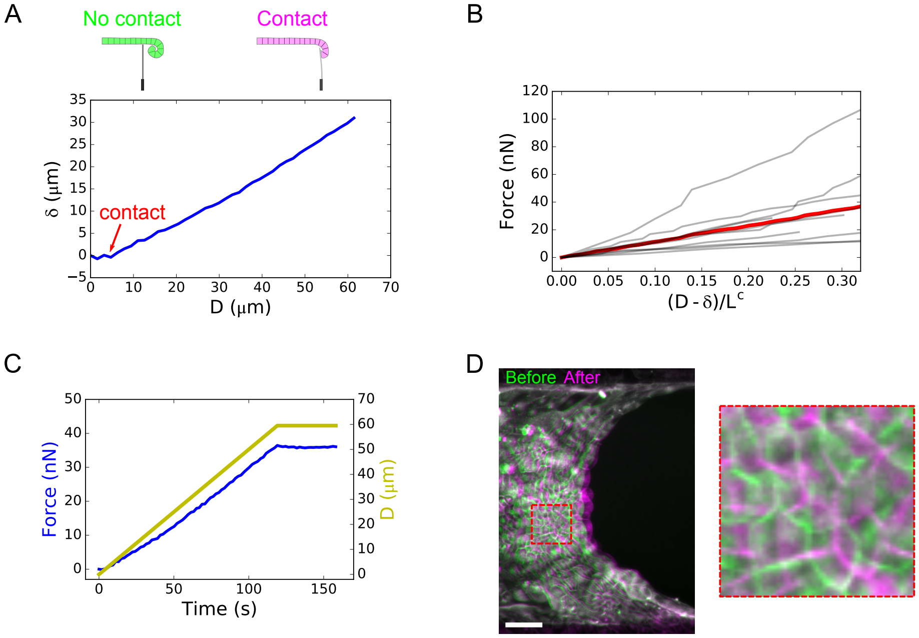


**Fig.S 3.** (A) Deflection $\delta$ of the cantilever during unfurling of a suspended MDCK monolayer as a function of the displacement $D$ imposed at its base by the motorized stage. The deflection is zero before contact with the tissue (see diagram above). The red arrow shows the contact point between the cantilever and the tissue. When the cantilever comes into contact with the curl and unfurls it, the restoring force of the curl results in a cantilever deflection. (B) Force variation in the cantilever during a ramp of displacement imposed at the base. The deformation of the tissue is represented by the ratio $\frac{D-\delta}{L^c}$, where $D - \delta$ is the displacement at the monolayer tip (see diagram in Figure 3A). The grey lines represent 9 individual datasets, the red line represents the average. (C) Representative graph of the temporal evolution of force in response to a ramp of displacement imposed at the cantilever base. Only negligible force change is observed after the movement is stopped, indicating that out-of-plane forces have an elastic origin and that a bending modulus of the tissue can be extracted from these measurements. (D) Overlay of scanning laser confocal images of the free edge of an MDCK monolayer (z-projection) before (green) and after (magenta) unfolding of the tissue with the force cantilever. The zoomed region (right) shows the displacement of cell outlines in the bulk of the monolayer. Cell membranes are marked with CellMask. Scale bar: 50μm.

### Figure S4

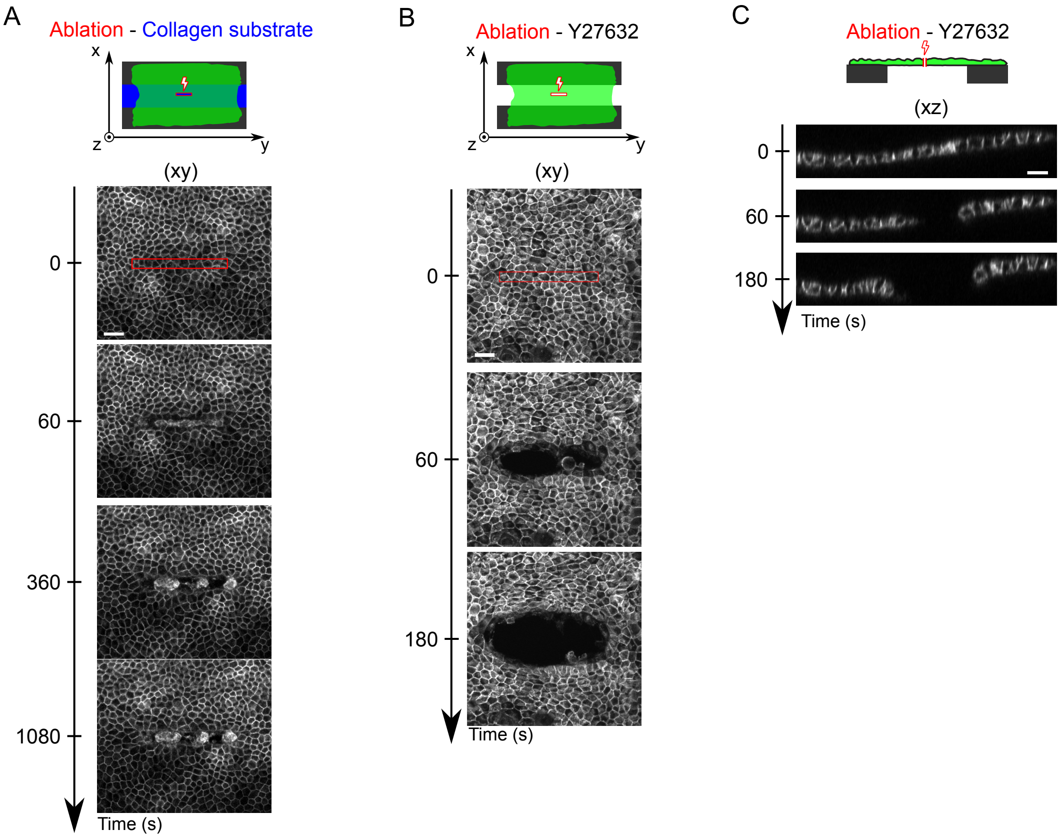


**Fig.S 4.** (A) Time series of ablation of an MDCK monolayer (green on the diagram) lying on a collagen substrate (blue) (N=6). The red rectangle indicates the ablated region. Laser ablation was performed after t=0s. Cell junctions are marked with E-Cadherin-GFP. Scale bar: 30μm. (B) Time series of the ablation of a suspended MDCK monolayer treated with 25μM Y27632, representative of N=10 monolayers. Laser ablation was performed in the frame after t=0s. Cell junctions are marked with E-Cadherin-GFP. Scale bar: 30μm. (C) Time-lapse profile view (xz-plane) of a suspended MDCK monolayer treated with 25μM Y27632 before (t=0s) and after laser ablation. Cell junctions are marked with E-Cadherin-GFP. Scale bar: 20μm.

### Figure S5

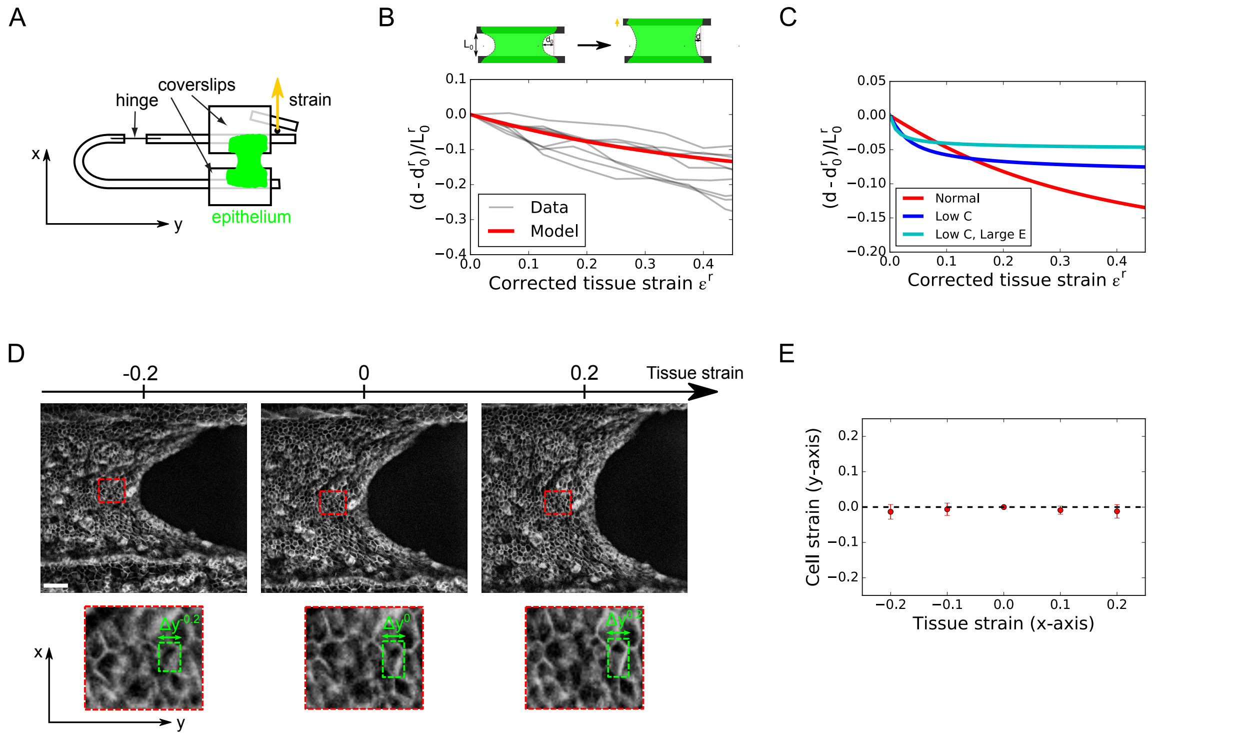


**Fig.S 5.** (A) Diagram showing the setup for application of uni-axial tissue-scale strain to suspended epithelia. (B) Variation of tissue deflection $d - d_0^r$ normalized by monolayer rest length $L_0^r$ as a function of in-plane corrected tissue strain $\epsilon^r = \frac{L-L_0^r}{L_0^r}$ . $d_0^r$ is the deflection at null strain. Here, tissue strain is corrected to have $L_0^r$ as the rest length of the epithelium. $L_0^r$ is defined experimentally as the deformation at which the tissue buckles under compression. It is inferior to $L_0$, which corresponds to the gap between coverslips after collagen digestion, because the tissue is naturally under tension after matrix removal (see (6)). The grey lines represent each experimental dataset, the red line represents the response predicted by the model with the average parameters measured in MDCK monolayers and no free parameters (see main manuscript and Appendix 2). (C) Same as A for the response predicted by the model with different parameters. The response with the average parameters measured in MDCK cells is shown in red (as in B). The response with reduced spontaneous curvature $\frac{1}{10}C_0^{MDCK}$ is shown in blue. The response with reduced spontaneous curvature and increased elastic modulus $10E^{MDCK}$ is shown in cyan. (D) Top: Confocal images of the free edge of a suspended MDCK monolayer for -20%, 0%, 20% tissue strain. Cell junctions are marked with E-Cadherin-GFP. Bottom: zoom on a region illustrating the negligible cell shape changes along the y-axis as cells are stretched and compressed along the x-axis ($\Delta y$ defines the width of the cell bounding box). Scale: 50µm. (E) Cell strain along the y-axis $\frac{\Delta y^\epsilon - \Delta y^0}{\Delta y^0}$ as a function of tissue stretch along the x-axis. Cells analyzed are chosen within the first five rows next to the curled region and close to where the monolayer edge is tangent to the x-axis (N=20 cells from 4 different monolayers).

### Figure S6

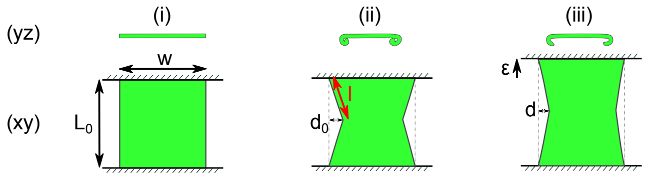


**Fig.S 6.** Schematic diagram showing the configurations of the elastic model used to determine the deflection of the monolayer free edge as a function of tissue in-plane strain. The monolayer is modeled as a rectangular sheet of length $L_0$ and width $w$ clamped on two of its sides. (i) Flattened configuration. (ii) Curled configuration. The free edge of length $2l$ is stretched and the monolayer curls giving rise to a deflection of the free edge, whose maximum is $d_0$ (iii) Curled and stretched configuration. The deflection of the free edge $d$ decreases in response to stretching (strain $\epsilon$).

#### Supplementary movies legend

**Movie 1.** Time-lapse of the free edge of an MDCK monolayer during the digestion of its collagen substrate by collagenase. Left: DIC imaging of the monolayer. Right: Collagen matrix imaged by confocal reflection microscopy. Collagenase is introduced in the medium at time t=0 min. Time is in min:sec. Scale bar: 50µm.

**Movie 2.** Time-lapse of the profile view of an MDCK monolayer during the digestion of its collagen substrate by collagenase. Collagenase is introduced in the medium at time t=0 min. Cell membranes are marked with CellMask. Time is in min:sec. Scale bar: 30µm.

**Movie 3.** Time-lapse of a Drosophila leg undergoing eversion. The outer peripodial membrane cracks and retracts, while the leg epithelium (center) changes shape. The red arrowhead indicates the retraction front. Cell junctions are marked with Armadillo-GFP (*Drosophila* homolog of β-catenin). Time is in min:sec. Scale bar: 50µm.

**Movie 4.** Time-lapse of a retracting peripodial epithelium imaged through Airyscan confocal microscopy. Top: Three-dimensional reconstruction. Bottom: Profile view perpendicular to the retraction front. Cell membranes are marked with CellMask. Time is in min:sec. Scale bar : 20µm.

**Movie 5.** Time lapse of an MDCK monolayer being unfurled by a cantilever that is displaced along the y-axis (profile view in the yz-plane). The medium is marked via Dextran-Alexa647 (red). Note that this dye also stains the tip of the cantilever underneath the monolayer, which facilitates the measurement of its displacement. Cell membranes are marked with CellMask (green). Time is in min:sec. Scale bar: 20µm.

**Movie 6.** Time-lapse of an MDCK monolayer grown on a collagen substrate undergoing laser ablation and subsequent wound healing. The red rectangle defines the border of the ablated region. Cell junctions are marked with E-Cadherin-GFP. Time is in min:sec. Scale bar: 30µm.

**Movie 7.** Time-lapse of a suspended MDCK monolayer undergoing laser ablation. The red rectangle defines the border of the ablated region. Cell junctions are marked with E-Cadherin-GFP. Time is in min:sec. Scale bar: 30µm.

**Movie 8.** Time-lapse of the profile of a suspended MDCK monolayer undergoing laser ablation. The profile view was taken along the plane perpendicular to the longest axis of the cut. Cell junctions (green) are marked with E-Cadherin-GFP, the medium (red) is marked with Dextran-Alexa647. Time is in min:sec. Scale bar: 20µm.

**Movie 9.** Time-lapse of the profile of a suspended MDCK monolayer treated with 25µM Y-27632 undergoing laser ablation. The profile view was taken along the plane perpendicular to the longest axis of the cut. Cell junctions are marked with E-Cadherin-GFP. Time is in min:sec. Scale bar: 20µm.

**Movie 10.** Time-lapse of the profile of a suspended MDCK monolayer before and after cut along the full length of the tissue at the interface with the coverslip. The profile view is taken along the plane perpendicular to the cut. Time is in min:sec. Scale bar: 30µm.

**Movie 11.** Time-lapse of the profile view of the free edge of a suspended MDCK monolayer during a ramp of in-plane strain. The profile view was taken along the plane perpendicular to the stretch axis. The movie starts at 0% strain, the monolayer is subsequently stretched to 90% strain, then compressed to -30% strain and finally brought back to 0%. (Strain is defined with reference to the monolayer length after digestion of collagen.) Cell membranes are marked with Cell Mask. Time is in min:sec. Scale bar: 30µm.

**Movie 12.** Time-lapse of the free edge of a suspended MDCK monolayer during a ramp of in-plane strain . The movie starts at 0% strain, the monolayer is subsequently stretched to 30% strain, compressed to -30% and brought back to 0%. (Strain is defined with reference to the monolayer length after digestion of collagen.) Time is in min:sec. Scale bar: 50µm.

**Movie 13.** Time-lapse of the profile of the free edge of a suspended MDCK monolayer before and after a step of in-plane strain (30% stretch). The profile view along the plane perpendicular to the stretch axis. Cell junctions are marked with E-Cadherin-GFP. Time is in min:sec. Scale bar: 30µm.
